# Maximum likelihood estimation of fitness components in experimental evolution

**DOI:** 10.1101/345660

**Authors:** Jingxian Liu, Jackson Champer, Chen Liu, Joan Chung, Riona Reeves, Anisha Luthra, Yoo Lim Lee, Andrew G. Clark, Philipp W. Messer

**Affiliations:** Department of Biological Statistics and Computational Biology; Department of Molecular Biology and Genetics, Cornell University, Ithaca, NY 14853

## Abstract

Estimating fitness differences between allelic variants is a central goal of experimental evolution. Current methods for inferring selection from allele frequency time series typically assume that evolutionary dynamics at the locus of interest can be described by a fixed selection coefficient. However, fitness is an aggregate of several components including mating success, fecundity, and viability, and distinguishing between these components could be critical in many scenarios. Here we develop a flexible maximum likelihood framework that can disentangle different components of fitness and estimate them individually in males and females from genotype frequency data. As a proof-of-principle, we apply our method to experimentally-evolved cage populations of *Drosophila melanogaster*, in which we tracked the relative frequencies of a loss-of-function and wild-type allele of *yellow*. This X-linked gene produces a recessive yellow phenotype when disrupted and is involved in male courtship ability. We find that the fitness costs of the yellow phenotype take the form of substantially reduced mating preference of wild-type females for yellow males, together with a modest reduction in the viability of yellow males and females. Our framework should be generally applicable to situations where it is important to quantify fitness components of specific genetic variants, including quantitative characterization of the population dynamics of CRISPR gene drives.

## Introduction

The concept of fitness lies at the core of Darwin’s theory of evolution by natural selection. If two allelic variants differ in fitness, the fitter allele should increase in population frequency over time at the expense of the less fit allele. The study of temporal changes in allele frequencies can therefore allow us to infer relative fitness differences. Various approaches have been developed to this end that can disentangle such selection from the stochastic frequency fluctuations generated by random genetic drift.^4, 15–18, 23–25, 27, 29, 35–37, 40–42^

The general principle of these methods is to devise a probabilistic model for expected allele frequency dynamics, often based on a Wright-Fisher process. Comparison of model predictions with empirical allele frequency measurements then allows inference of model parameters (which may include effective population size) using statistical approaches such as maximum likelihood (ML) or Bayesian inference. Applications of these methods have provided key insights into the strength of selection operating on particular alleles.^26, 30^

A common assumption in these methods is that selection between two allelic variants can be described by a single selection coefficient. While often reasonable, it will be critical in many scenarios to distinguish among the individual components that constitute fitness.^9, 33, 39^ For example, preferential mate choice cannot typically be mapped onto a logistic growth model specified by a fixed selection coefficient, since it could result in frequency-dependent dynamics.^2^ Individual fitness effects could also often differ between males and females, which would lead to systematic biases in genotype frequencies between sexes when selection is sufficiently strong.

Rigorous inference of selection can include four major selection components: zygotic selection (viability from zygote to adult), sexual selection, fecundity selection, and gametic selection that usually involves genetic elements with biased inheritance.^9, 33^ Evaluation of these four selection components in a natural population requires detailed population monitoring, as observations need to be recorded at four different life-cycle stages.^9, 33, 34, 38^ Typically, the analysis of selection components in the laboratory involves performing a series of isolated experiments designed to individually quantify each component.^5^ For example, progeny counts from controlled crosses and backcrosses can reveal differences in zygote-to-adult viability as measured by deviation from Mendelian proportions.^32^ Egg counts from controlled crosses can reveal genotype-specific differences in fecundity. Mating arenas with either observed matings or subsequent scoring of progeny can allow for estimation of sexual selection.^12^ The challenge with all of these methods is that it is extremely difficult to know if all attributes of a gene that may result in differential propagation have been considered, although it is possible to test the correspondence between individual fitness components estimates and allele frequency dynamics in cage populations.^10, 11^

In this study we develop a ML inference framework that can disentangle sexual, fecundity, and viability selection from genotype frequency time series data. Our analytical approach employs a continuous extension of the multinomial distribution, allowing us to infer effective population size simultaneously with selection parameters. As a proof-of-principle, we apply our inference framework to empirical data obtained from cage evolution experiments of *Drosophila melanogaster*, in which we tracked relative genotype frequencies of a wild-type and mutant version of *yellow*, an X-linked gene required for black pigment formation.^14^ The mutant allele disrupts *yellow*, resulting in a recessive *yellow* body phenotype. The *yellow* gene is also required for the wing extension behavior that is part of male courtship in *D. melanogaster*.^3^,^44^ Consequently, disruption of *yellow* affects mating competitiveness in male flies.^13, 21^

We find that in cage experiments, our ML inference method can robustly distinguish between several selection models involving preferential mate choice, fecundity, and viability, and further distinguish between male and female fitness costs. Such measurements should become particularly important in scenarios where fitness costs are large and allele frequencies are expected to change rapidly, as will likely be the case for recently proposed CRISPR gene drive approaches.^6^

## Methods

### Plasmid construct design and generation of transgenic fly line

We designed a construct targeting the X-linked *yellow* gene in *D. melanogaster*. Disruption of this gene causes a recessive yellow phenotype, specified by a lack of dark pigment in the adult cuticle. Our construct additionally encodes a dsRed protein driven by a 3xP3 promoter, which produces an easily identifiable fluorescent phenotype. dsRed is not visible in wild-type eyes in the Canton-S background, but we observed dsRed expression in the abdomen of younger insects and in the ocelli.

The donor plasmid BHDyR was constructed by Gibson assembly of the restriction digest of IHDyV1^8^ with StuI and XhoI and PCR amplification of the pDsRed-attp plasmid^20^ with the oligos dsRedY_F: GGGTT-TTGGACACTGGGAATTCTTGCATGGCTAGACGAAGTTATCGTACGGGATCTAAT and dsRedY_R:TTAGTGGTGGTATTGCCGATGCCCACGGACGCGCCGGTTAAGATACATTGATGAGTTTGG (IDT). Gibson assembly of plasmids was performed with Assembly Master Mix (New England Biolabs), and plasmids were transformed into JM109 competent cells (Zymo Research).

The transgenic line in the study was transformed at GenetiVision by injecting the BHDyR donor plasmid into Canton-S *D. melanogaster* embryos. Cas9 was provided by co-injection with plasmid pHsp70-Cas9^19^ (a gift from Melissa Harrison, Kate O’Connor-Giles, and Jill Wildonger, Addgene plasmid 45945), and a gRNA plasmid (BHDyg1)^8^ was also included in the injection. Concentrations of donor, Cas9, and gRNA plasmids were 98, 94, and 58 ng/*μ*L, respectively, in 10 mM Tris-HCl, 23 *μ*M EDTA, pH 8.1 solution. The injected plasmids were purified with ZymoPure Midiprep kit (Zymo Research). To obtain a homozygous fly line, the injected embryos were reared and crossed with wild-type Canton-S flies. The progeny with dsRed fluorescent protein in the eyes and abdomen, which indicated successful insertion of the construct, were selected and crossed with each other. The stock was considered homozygous when all male progeny had dsRed fluorescence for two consecutive generations.

### Fly rearing and phenotyping

Flies were reared at 25 ° C with a 14/10 hr day/night cycle. Bloomington Standard medium was provided as food every two weeks. During initial phenotyping, flies were anesthetized with CO_2_ and examined with a stereo dissecting microscope, and the fluorescent red phenotype was observed using the NIGHTSEA system.

For cage studies, enclosures of internal dimensions 30×30×30 cm (Bugdorm, BD43030D) were used to house flies. At the start of an experiment, transgenic and Canton-S flies of approximately the same age were separately allowed to lay eggs in food bottles for two days. These food bottles (nine for cage 1, eight for cage 2, five for Cages 3-5) were then placed in cages at the desired starting ratio between transgenics and Canton-S flies. Eleven days later, bottles were replaced in the cage with fresh food (at a 1:1 ratio for cage 1 and 2:1 for other cages), leaving adult flies. Two days later, bottles were removed again from the cages and flies retained, and fresh food bottles were added (nine for cage 1, eight for cage 2, five for cages 3-5). One day later, flies were frozen for later phenotyping, and food bottles were retained for eleven days to allow the next generation to hatch. This cycle was repeated until the completion of the experiments.

## Results

### Cage evolution experiments

We created a transgenic *D. melanogaster* line based on the Canton-S strain, in which we inserted a construct expressing a dsRed fluorescent protein into the *yellow* gene. This insertion effectively disrupts the gene, producing the characteristic yellow phenotype, which is recessive. At the same time, expression of the dsRed protein produces an easily identifiable, co-dominant phenotype characterized by red fluorescence in the eyes, abdomen, and ocelli, though only the latter two could be observed in Canton-S flies due to eye pigmentation.

We will refer to the allele with the inserted construct as *y*, while we will denote the wild-type allele as +. Because *yellow* is located on the X chromosome, males of genotype *y* are expected to show both the yellow (body) and red (eye) phenotypes, whereas + males are expected to show neither phenotype. In females, *yy* homozygotes are expected to show both the yellow and red phenotypes, *y*+ heterozygotes are expected to show only the red phenotype, and ++ homozygotes are expected to show neither phenotype. Our system thereby allows us to unambiguously ascertain genotypes at the *yellow* locus from phenotypic assays of individual flies.

We evolved five laboratory cage populations, each initialized from a mixture of wild-type Canton-S flies and our transgenic *yellow* flies. Genotype frequencies at the yellow locus were then tracked over the course of several generations. Our first two cages were evolved over six (cage 1) and five (cage 2) generations, starting with initial *y* allele frequencies of ~70%. Three additional cages were each evolved for just a single generation with starting frequencies of the *y* allele of ~20% for cage 3, ~50% for cage 4, and ~80% for cage 5. Each cage comprised between several hundred and several thousand flies. The observed genotype frequency trajectories and changes in male and female population size over time are shown in Figure 1.

**Figure 1:**
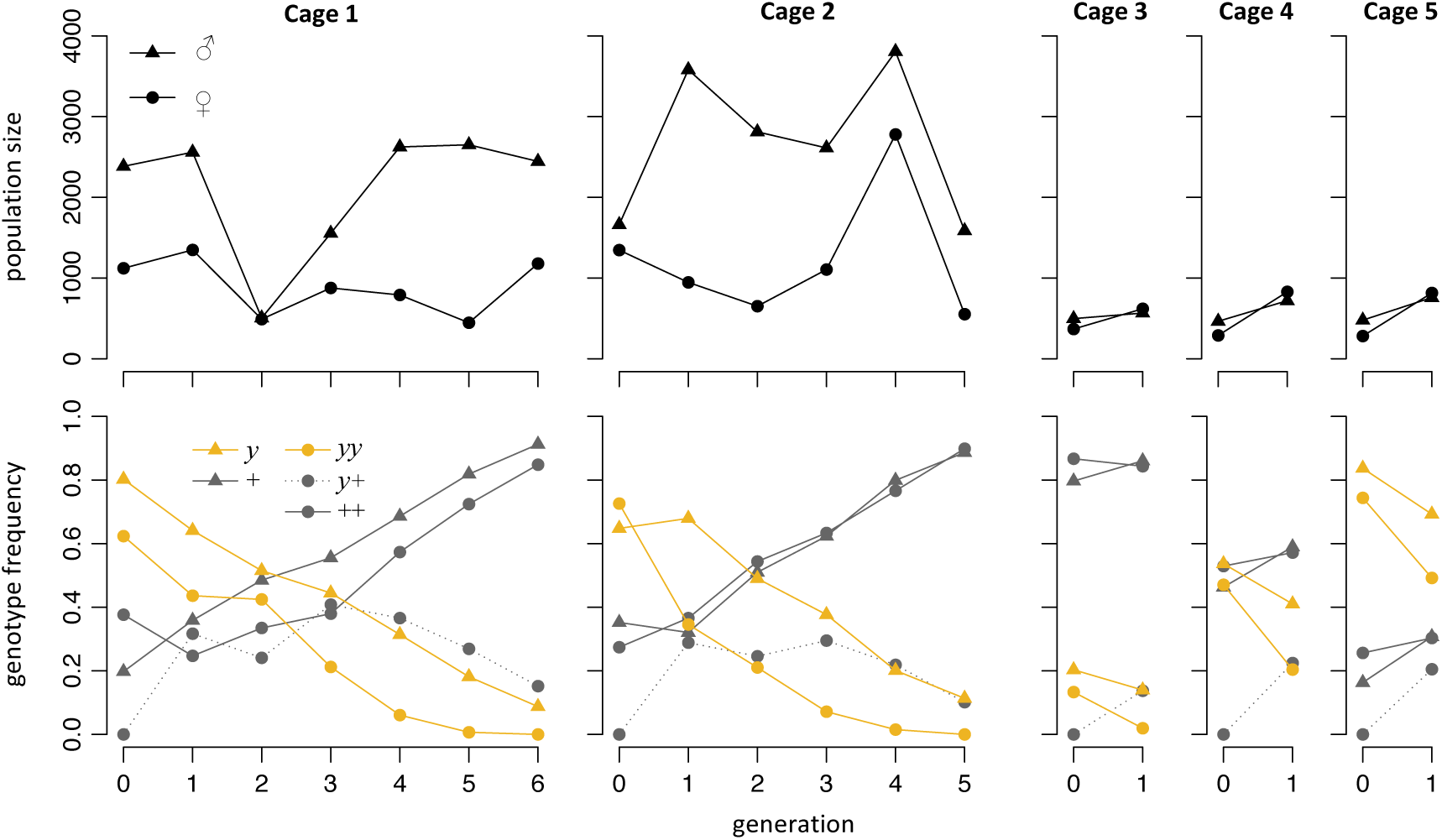
Population sizes and genotype frequencies in our five cage experiments. Note that we define genotype frequencies relatively among males, and relatively among females.

The frequency of the *y* allele decreased systematically in all five cages, confirming that *yellow* disruption is likely associated with a substantial fitness cost. In cage 1, for example, the *y* allele decreased from ~70% to less than 10% over the course of just six generations, and a broadly consistent rate of decline was observed among the other cages as well.

Census population sizes fluctuated noticeably in our cages. However, we do not believe that these fluctuations are the result of selection at the *yellow* locus, but rather reflect unrelated factors, such as larval competition and variance in food preparation. We also usually observed a bias in sex ratio of phenotyped flies since females had higher mortality from laying eggs in the crowded food bottles, while males approached the food bottles less often.

### Evolutionary model

The observation of rapidly declining *y*-allele frequency in all five population cages raises the possibility that we may have statistical power to estimate individual fitness components associated with *yellow* disruption. To explore this possibility, we will first establish an evolutionary model for allele frequency dynamics in our experiments that incorporates three specific components of fitness: viability, fecundity, and mating success.

Consider an X-linked locus with two segregating alleles: wild-type (+) and yellow (*y*). The *y* allele produces the yellow phenotype, which we assume has lower fitness than the wild-type phenotype. These fitness costs could be due to reduced viability, fecundity, and/or mating success in individuals with yellow phenotype compared to wild-type individuals. In the following, we will describe how each of these three different fitness components is incorporated into our model. In all cases, we assume that the *y* allele is strictly recessive, so that wild-type and *y*+ heterozygous females have the same fitness. We will denote the counts of individuals observed in the present generation *t*, partitioned by genotype, as *N_y_*, *N_+_*, *N_yy_*, *N_y_*_+_, *N*_++_ (the first two are males, the last three are females).

Viability specifies the probability that a zygote survives to reproductive age, which we assume will depend only on the phenotype and sex of the individual. Specifically, we define *v_m_* to be the relative viability of yellow males compared to wild-type males, and *v_f_* to be the relative viability of yellow females compared to wild-type females:

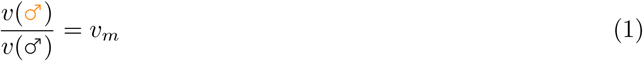

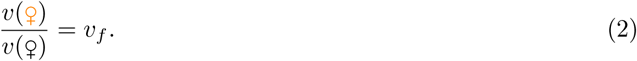

Using relative viabilities among males and females, rather than absolute viabilities, can help prevent systematic biases due to the unequal sex ratios in our cages.

Fecundity specifies the reproductive success of a mating pair, as measured by the expected number of offspring. In contrast to viability, which is a function of an individual zygote, fecundity is therefore a function of a mother-father pair. In our model, we assume that fecundities are defined by the following mating table, specifying the relative fecundities of matings involving parents with yellow phenotype, compared to matings of wild-type mothers with wild-type fathers:

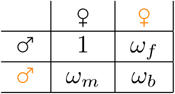

For our sexual selection model, we assume a scenario of female mate choice, in which females will choose mates with a potential preference based on phenotype. We define the actual probabilities that a female of a particular phenotype chooses a male of a particular phenotype to be:

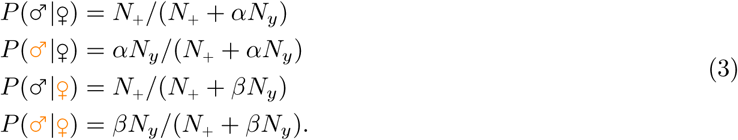

Here the parameters *α* and *β* specify the relative reduction in mating success of yellow males compared to wild-type males in mating with wild-type or yellow females, respectively.

Biologically there is no reason to assume that yellow phenotype males or females would actually have higher viability, fecundity, or mating success than their wild-type counterparts. Thus, we will assume that all of our selection parameters are constrained within 0 ≤ *v_m_*, *v_f_*, *ω_m_*, *ω_f_*, *ω_b_*, *α*, *β* ≤ 1, which will also help prevent overfitting in our ML inference approach. Finally, we assume discrete generations and random segregation of alleles within gametes due to a lack of any known mechanism for biased inheritance of *y*. Figure (2) illustrates the life cycle in this evolutionary model over the course of one generation, depicting where the different types of selection operate.

**Figure 2:**
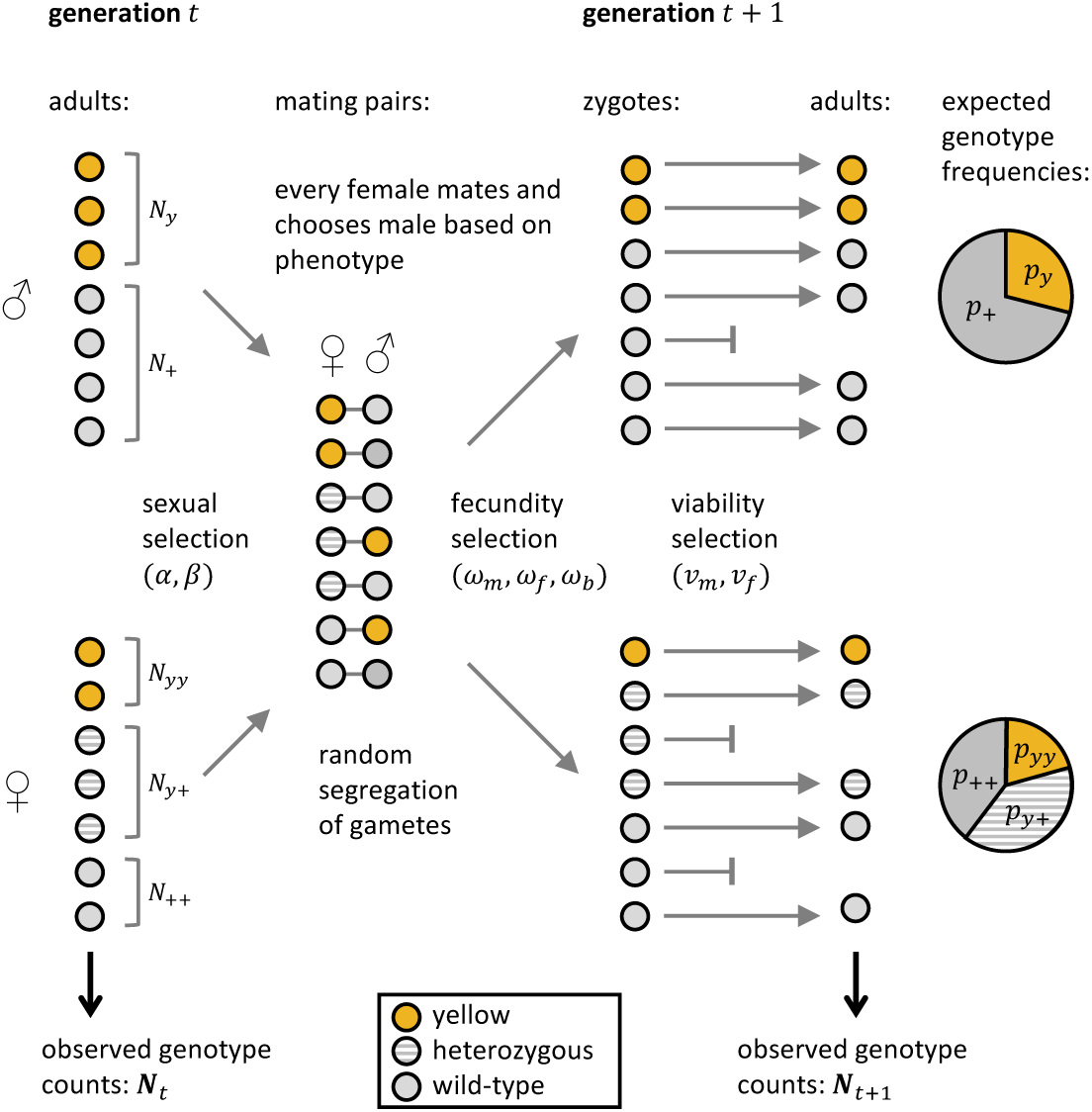
Illustration of the different stages of the life cycle during one generation in our model.

With these definitions at hand, we can calculate the expected values of genotype frequencies in generation *t*+1 as a function of their frequencies in generation *t* and the parameters of the evolutionary model. At our X-linked locus, male zygotes always inherit their X chromosomes from their mothers, while females inherit one copy from each parent. When assuming absence of any selection (*v_m_* = *v_f_* = *ω_m_* = *ω_f_* = *ω_b_* = *α* = *β* = 1), the expected genotype frequencies in the next generation will then be given by:

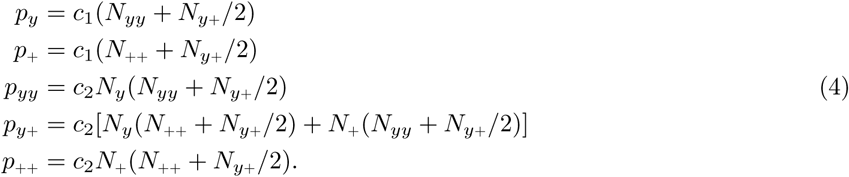

Here *p_y_* and *p*_+_ denote the expected relative frequencies of *y* and + genotypes among males, and *p_yy_*, *p_y_*_+_, and *p*_++_ denote the expected relative frequencies of *yy*, *y*+, and ++ genotypes among females. The normalization coefficients *c*_1_ and *c*_2_ are defined by the conditions that gamete frequencies in males and females each have to sum to one: *p_y_* + *p*_+_ = *p_yy_* + *py*_+_ + *p*_++_ = 1. Note that we do not use absolute genotype frequencies in the population, but only relative frequencies in males and females, again to prevent any systematic biases due to unequal in sex ratios in our cages.

Viability selection is straightforward to incorporate into this framework: we can simply interpret the genotype frequencies derived in (4) as the expected frequencies of zygotes, which then just need to be multiplied by the corresponding viability coefficients (and re-normalized). To incorporate fecundity and mate choice, however, we also have to incorporate the specific probabilities that each mating that produces the respective zygote genotype actually occurs, and then weigh them by their fecundities. Altogether, this yields our full model:

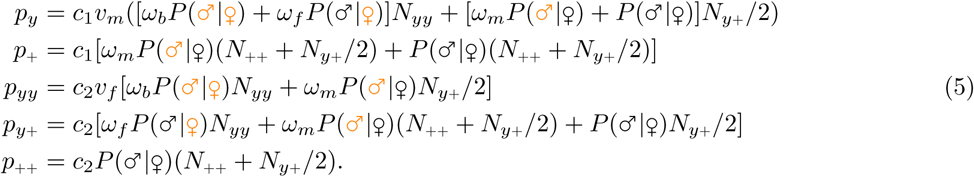

The coefficients *c*_1_ and *c*_2_ are again defined by normalization. Note that although this model is specific for an X-linked gene, it is in fact easy to extend to other inheritance patterns, such as an autosomal locus. The general principle is always the same: to calculate the expected frequency of a given genotype, we have to sum over all matings that can produce this genotype, weight each by its respective mating probability and expected fecundity of the mating pair, then multiply all genotypes by their respective viabilities, and finally normalize all genotype frequencies. Other selection scenarios, such as different dominance effects, could be easily implemented as well by changing the genotypes to which fitness parameters are applied.

### ML inference framework

The evolutionary model developed above allows us to calculate the expected changes in genotype frequencies across generations for a given set of model parameters. In turn, we may be able to use these relations for inferring the model parameters, given observed genotype frequency changes. To perform this inference in a ML framework, we require a probabilistic model that can specify the actual probabilities of observing a given set of genotype frequencies in the next generation, which will fluctuate around their expected values because of random genetic drift.

We will formally describe these fluctuations using a diploid Wright-Fisher model, where each individual in generation *t* + 1 has its genotype sampled randomly from a multinomial probability distribution, specified by the expected genotype frequencies from Eq. (5) and an effective population size (*N_e_*) controlling the level of random genetic drift in the population. Note that we expect the effective population size of our cages to be considerably smaller than their census sizes due to a variety of factors, such as a higher variance in offspring numbers among the flies in our cages (especially males) compared to a Wright-Fisher population.

There is an important complication when it comes to calculating probabilities in this model: In a Wright-Fisher population of effective size *N_e_*, the probability of observing a given vector 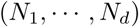 of counts of all *d* possible genotypes is specified by a multinomial distribution, which is properly defined only when 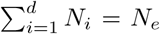. However, this condition will typically be violated in our cages, as we expect sample sizes (constituted by all individuals present in a cage in our experiments) to be much larger than *N_e_*.

To address this problem, we will employ a continuous approximation for the multinomial distribution, where genotype counts are replaced by genotype frequencies and the discrete multinomial probabilities are replaced by a multi-dimensional probability density. We define this density by using the gamma function Γ(*x* + 1) = *x*! as a natural continuous extension for the factorials in the multinomial distribution. Specifically, consider a diploid Wright-Fisher model with effective population size *N_e_* and *d* possible genotypes, where 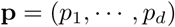 specifies their expected frequencies in generation *t* + 1. The probability density of observing a given set of genotype frequencies 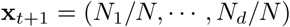 in generation *t* + 1 is then defined by:

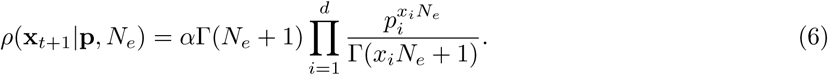

The constant *α* is to ensure normalization when *ρ* is integrated over the *d*-dimensional standard simplex defined by the constraint ∑*_i_ x_i_* = 1. In this framework, the probability of observing genotype frequencies that fall inside a given area Ω of the *d*-dimensional frequency space is then obtained by integrating the probability density over this area: Pr(x ∊ Ω) = ∫_Ω_ *ρ*(x)dx.

Unfortunately we are not aware of any closed-form analytic expressions for the normalization coefficient α, except for the special case *d* = 2, but we can derive a workable approximation by discrete partitioning. Specifically, we will count the number of possible frequency vectors assuming that frequencies *x_i_* can still only take on discrete values 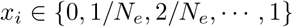 on the simplex ∑*_i_ x_i_* = 1. This number is given by the binomial coefficient

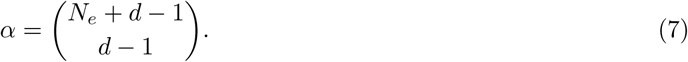

Note that under this choice of α, the function *ρ*(**x**) is technically no longer a proper probability density, as integration over the continuous frequency simplex does not always yield a value of exactly one. This is because our approximation of *α* is still based on a discrete partitioning of the frequency space, which should converge to the continuous limit only as *N_e_* goes to infinity. However, these deviations become noticeable primarily for very small values of *N_e_*, where the applicability of the Wright-Fisher model becomes questionable in general. Our simulations below also confirm that this discrete approximation works well in practice.

To allow for the possibility of biased sex ratios in our model, while avoiding specific assumptions about what caused these biases in our cages, we will further factorize the overall probability density of observed genotype counts into the individual probabilities of observing the male and female genotype counts independently:

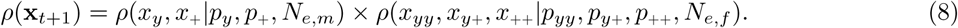

Here *x_y_* and *x*_+_ specify the observed genotype frequencies in generation *t* + 1 in males, and *x_yy_*, *x_y_*_+_, *x*_++_ specify the observed genotype frequencies in females. For males, we have *d* = 2, and the normalization factor in *ρ* is therefore *α* = (*N_e,m_* + 1). For females, we have *d* = 3, and therefore *α* = (*N_e_*_,*f*_ + 2)(*N_e_*_,*f*_ + 1)/2.

The factorization employed in definition (8) provides a very flexible framework for defining effective population size, which could be assumed to differ between males and females or could vary based on the census population size. We will discuss this issue in more detail below. For now, we want to assume the simplest model of a constant effective population size with equal sex ratio:

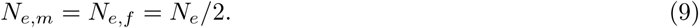

With these definitions in place, we can formulate the likelihood density function for the parameter vector *θ* = {*N_e_*, *v_m_*, *v_f_*, *ω_m_*, *ω_f_*_,_ *ω_b_*_,_ *α*, *β*}, given the vectors of observed genotype counts in generations *t* and *t* + 1 of a population cage. Multiplying these individual likelihoods across all generations 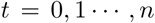 in an experiment (i.e., assuming independence across generations) then yields the log-likelihood density function:

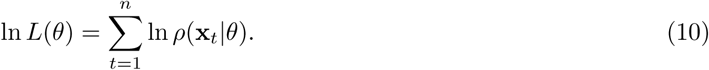

We can easily extend this approach across multiple cages by simply summing the log-likelihoods of the individual cages. The resulting log-likelihood density function can then be used for inference of maximum likelihood estimates (MLEs) of the parameter values and their confidence intervals.

Several aspects of the above approach are worth further mention: First, *N_e_* has now become just another parameter of the model and can thus be inferred the same way as we do with the other parameters. This way, *N_e_* can also take on continuous values in our model. Second, our ML inference approach is very general and could be easily applied to other evolutionary models as well, so long as the model provides expected values for genotype frequencies, given their frequencies in the previous generation and model parameters. Finally, we want to acknowledge that we have not explicitly incorporated sampling variance into our framework, since we assume that the observed genotype counts reflect the actual numbers of individuals that were present in a cage. While some sampling noise could of course have occurred in any given generation of an experiment, we expect that the resulting variance in genotype frequencies should be negligible compared to the variance due to drift. This is also supported by the fact that we typically infer effective population sizes that are an order of magnitude smaller than the census sizes of our cages.

### Power evaluation

Even when combined over all five cages, the genotype trajectories in our experiments comprise a total of only 14 transitions between consecutive generations. In light of such limited data, can we still expect our ML framework to have sufficient power to infer the different parameters of our model? To explore this question and also provide a general proof-of-principle of our inference framework, we first tested it on simulated data.

Figure 3A shows the outcomes of three Wright-Fisher simulation runs, in which we modeled genotype frequency trajectories over 10 generations in a cage under three different selection scenarios, differing in whether fitness costs of an X-linked allele were due to reduced mating success, reduced viability, or reduced fecundity. These initial simulations were run without drift (in the limit of “infinite” *N_e_*) to assess whether we can detect systematic differences between the three scenarios. We tuned the selection strength in each model to roughly conform with the observed decay-rate of *y* allele frequencies in our experimental cages.

**Figure 3:**
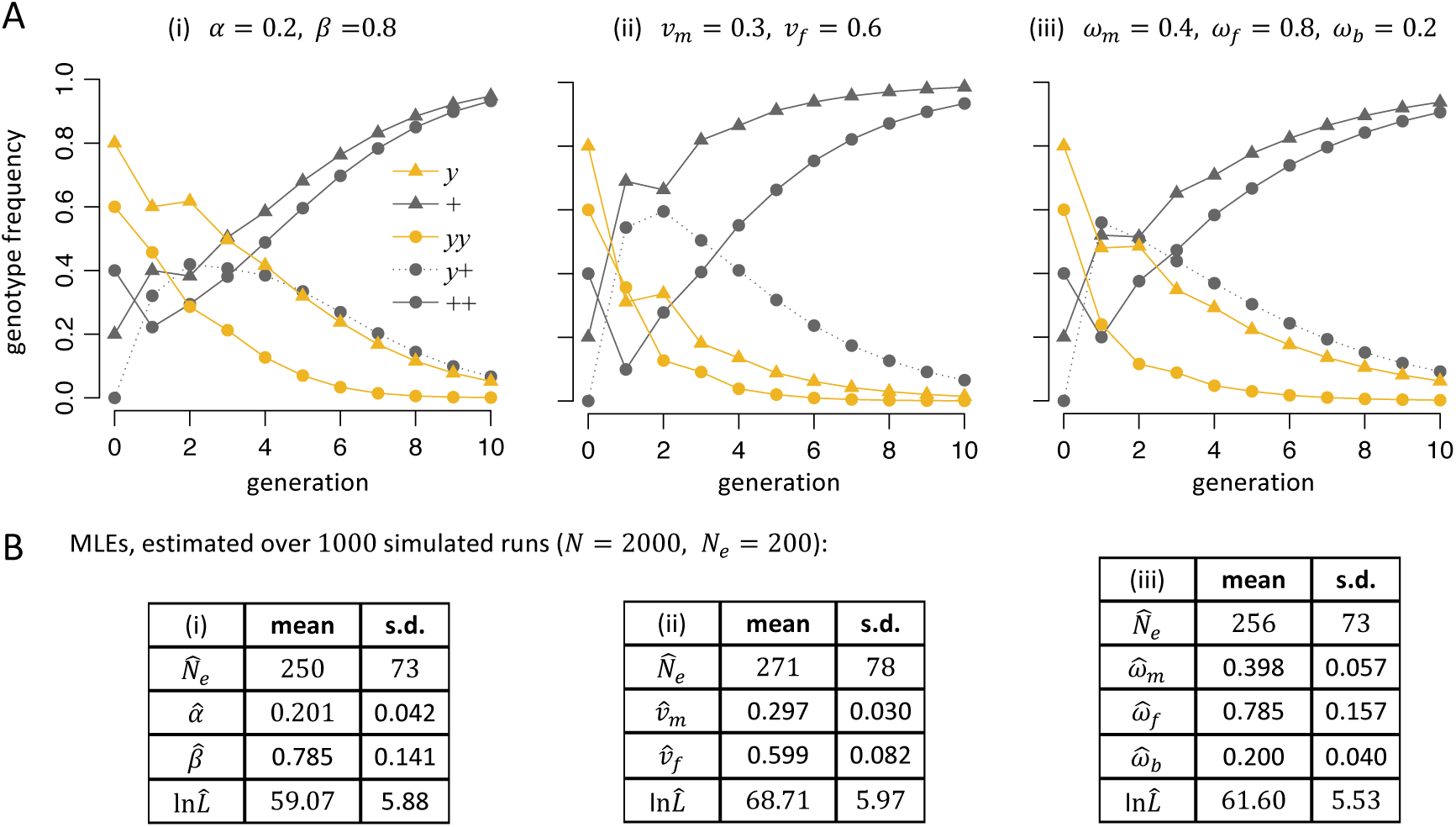
(A) Simulated genotype frequency trajectories under different selection scenarios in a deterministic model without drift: (i) sexual selection only (*α* = 0.2, *β* = 0.8; *v_m_* = *v_f_* = *ω_m_* = *ω_f_* = *ω_b_* = 1); (ii) viability selection only (*v_m_* = 0.3, *v_f_* = 0.6; *ω_m_* = *ω_f_* = *ω_b_* = *α* = *β* = 1); (iii) fecundity selection only (*ω_m_* = 0.4, *ω_f_* = 0.8, *ω_b_* = 0.2; *v_m_* = *v_f_* = *α* = *β* = 1). (B) Accuracy of our ML inference framework for the three selection scenarios. Tables show the average parameter MLEs and standard deviations estimated over 1000 simulation runs of the specific scenario. These simulations were run with drift, assuming populations of 2000 individuals with *N_e_* = 200. For each scenario, inference was performed by only inferring the parameters of the given model, while keeping the other parameters fixed at one. In all three scenarios, the parameter MLEs converge to their true values when averaging over individual runs.

The simulations reveal that there are indeed systematic differences in the expected genotype frequency trajectories between the three scenarios, even though the overall reduction in *y*-allele frequencies after 10 generations were similar. While differences are most pronounced in the early generations of a run, where frequency changes are the largest, they also extend throughout the run, as can be seen in the different frequency ratios of *y* males and *yy* females. These results suggest that, at least in principle, different fitness components could be disentangled by our approach.

Figure 3B demonstrates that even in a model with substantial drift (*N_e_* = 200), our ML approach can still infer the individual model parameters from a single 10-generation experiment if selection is sufficiently strong. The average MLEs for the selection parameters converge closely to the true values, with standard deviations among individual simulation runs typically on the order of 10 − 15% of the parameter values. However, the *N_e_* estimates are consistently higher then the simulated values. This is likely due to the fact that in any single experiment, drift alone should already result in an upward or downward shift in yellow allele frequency, which our method would likely infer as “additional” selection, thereby underestimating the true amount of drift. Consider, for example, a completely neutral scenario. In this case, our method would still likely infer some positive or negative selection in any given run, leading to systematically lower inferred levels of drift, even though the direction of selection would be random and average out over many runs.

So far, we have only tested our inference method under the assumption that we know which type of selection has acted. In this case, we validated that the method can reliably infer the respective selection parameters, but can it also distinguish the different types of selection from each other? To test this, we applied our method to each of the three selection scenarios, using either the correct inference model or models in which a different type of selection was assumed. This can be easily implemented in our framework by fixing specific selection parameters. For example, to devise a sexual-selection-only inference model, the parameters *α* and *β* would be allowed to vary, while all other selection parameters would be set to one. The resulting parameter MLEs and maximum log-likelihoods values for each simulation model / inference model combination are shown in Table 1. As expected, the highest log-likelihoods were always achieved when the simulation model matched the inference model, and the decreases in likelihood when the inference model was misspecified were significant in all cases (*p* < 0.05) according to likelihood ratio tests. This indicates that at least for situations when model parameters have distinctly different values, we should have sufficient power to determine the type of selection operating on a particular system.

**Table 1:** Disentangling sexual, viability, and fecundity selection. The columns specify the three simulation models from Fig. 3. The rows specify three different inference models, assuming either a model of sexual selection only (free parameters *α*, *β*), viability selection only (free parameters *v_m_*, *v_f_*), or fecundity selection only (free parameters *ω_m_*,*ω_f_*, *ω_b_*). The cells show the resulting parameter MLEs and maximum log-likelihood values when the particular inference model is applied to the particular simulation model, averaged over 1000 simulation runs in each case.

|  |  | Simulated model: |  |  |
| --- | --- | --- | --- | --- |
| | | (i) sexual<br>sel. only<br>$\alpha = 0.2$<br>$\beta = 0.8$ | (ii) viability<br>sel. only<br>$v_m = 0.3$<br>$v_f = 0.6$ | (iii) fecundity<br>sel. only<br>$\omega_m = 0.4$<br>$\omega_f = 0.8$<br>$\omega_b = 0.2$ |
| Model used for inference: | sexual<br>sel. only | $\ln \hat{L} = 59.1$<br>$\hat{\alpha} = 0.20$<br>$\hat{\beta} = 0.79$ | $\ln \hat{L} = 34.1$<br>$\hat{\alpha} = 0.78$<br>$\hat{\beta} = 0.37$ | $\ln \hat{L} = 56.7$<br>$\hat{\alpha} = 0.35$<br>$\hat{\beta} = 0.20$ |
| | viability<br>sel. only | $\ln \hat{L} = 46.2$<br>$\hat{v}_m = 0.97$<br>$\hat{v}_f = 0.65$ | $\ln \hat{L} = 68.7$<br>$\hat{v}_m = 0.30$<br>$\hat{v}_f = 0.60$ | $\ln \hat{L} = 45.9$<br>$\hat{v}_m = 0.87$<br>$\hat{v}_f = 0.39$ |
| | fecundity<br>sel. only | $\ln \hat{L} = 58.0$<br>$\hat{\omega}_m = 0.24$<br>$\hat{\omega}_f = 0.58$<br>$\hat{\omega}_b = 0.42$ | $\ln \hat{L} = 42.7$<br>$\hat{\omega}_m = 0.96$<br>$\hat{\omega}_f = 0.02$<br>$\hat{\omega}_b = 0.35$ | $\ln \hat{L} = 61.6$<br>$\hat{\omega}_m = 0.40$<br>$\hat{\omega}_f = 0.79$<br>$\hat{\omega}_b = 0.20$ |

### Parameter estimation in cage experiments and model comparison

We applied our ML inference method to the combined data from our five cage populations by adding the individual log-likelihoods of each of the 14 generational transitions observed in our experiments. Table 2 shows the resulting parameter MLEs and maximum log-likelihood values for several different inference models.

**Table 2:** Parameter MLEs and model comparison. Each row shows the parameter MLEs and associated maximum log-likelihood value when estimated from the combined data across all cages (1-5). An entry of 1* indicates that the parameter was fixed at a value of one in the particular inference model. “=” indicates that the value of a parameter is fixed in relation to another. *p* denotes the number of free parameters of the model, and AICc specifies its corrected Akaike information criterion value, 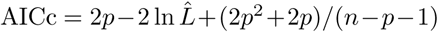, where *n* = 14 is the sample size. Smaller AICc values indicate better support for the model.

| model description | $\hat{N}_e$ | $\hat{v}_m$ | $\hat{v}_f$ | $\hat{\omega}_m$ | $\hat{\omega}_f$ | $\hat{\omega}_b$ | $\hat{\alpha}$ | $\hat{\beta}$ | $\ln \hat{L}$ | $p$ | AICc |
| --- | --- | --- | --- | --- | --- | --- | --- | --- | --- | --- | --- |
| full model | 167 | 0.84 | 0.76 | 0.98 | 0.93 | 1.00 | 0.07 | 1.00 | 69.1 | 8 | -93 |
| no sexual sel. | 104 | 0.99 | 1.00 | 0.11 | 0.39 | 0.29 | 1* | 1* | 59.1 | 6 | -94 |
| no viability sel. | 156 | 1* | 1* | 0.82 | 0.87 | 0.73 | 0.09 | 1.00 | 67.7 | 6 | -111 |
| no fecundity sel. | 166 | 0.83 | 0.78 | 1* | 1* | 1* | 0.06 | 1.00 | 69.1 | 5 | -121 |
| sexual sel. only | 134 | 1* | 1* | 1* | 1* | 1* | 0.03 | 0.71 | 64.5 | 3 | -121 |
| viability sel. only | 16 | 0.83 | 0.62 | 1* | 1* | 1* | 1* | 1* | 16.0 | 3 | -24 |
| fecundity sel. only | 104 | 1* | 1* | 0.11 | 0.39 | 0.29 | 1* | 1* | 59.1 | 4 | -106 |
| simplistic model | 31 | 1* | 1* | 0.50 | $= \omega_m$ | $= \omega_m^2$ | 1* | 1* | 34.6 | 2 | -64 |
| minimal model 1 | 165 | 0.81 | $= v_m$ | 1* | 1* | 1* | 0.06 | 1* | 68.9 | 3 | -129 |
| minimal model 2 | 156 | 1* | 1* | 0.86 | $= \omega_m$ | $= \omega_m^2$ | 0.09 | 1* | 67.7 | 3 | -127 |

We first tested our full inference model with all three types of selection possible (eight free parameters total, including *N_e_*). The resulting parameter MLEs for this model suggest that all three types of selection were indeed acting in the cages, as indicated by the fact that at least one parameter for each type of selection was inferred to be different from one. However, we suspected that there could still be substantial overfitting in this model, given the rather large number of parameters for the amount of data available. Furthermore, various parameters could be intertwined with each other and difficult to disentangle in practice. For example, it should generally be possible to approximate many scenarios of referential mate choice to some extent by appropriately tuning a model with only fecundity selection, especially since the latter has more free parameters in our framework.

To assess which fitness components are indeed most essential to incorporate for accurately describing the genotype dynamics observed in our cages, we tested several inference models, systematically reducing model complexity. The quality of these models was then compared by calculating the corrected Akaike information criterion (AICc) values,^1, 22^ which provide an estimator for the goodness-of-fit of a given model while also penalizing an increase in the number of model parameters. While the correction term is strictly accurate only for linear models, it serves as an adequate approximation for more complex scenarios in most cases.^22^

We first evaluated inference models in which one type of selection was completely eliminated (by setting all parameters describing that component to a value of one). In this case, the largest reduction in maximum log-likelihood was observed when sexual selection was excluded. Inference models without viability or fecundity selection also had decreased maximum log-likelihoods compared to the full model, but reductions were less dramatic. All of these models yielded better AICc values than the full model because of their reduced number of parameters.

We next tested models with only one type of selection acting. Consistent with the above result, we found that a model with only sexual selection achieved a higher maximum log-likelihood compared to models with only viability or only fecundity selection, and the AICc value of this model was again better than the full model. The model with only viability selection was the least-supported model we tested overall.

A simplistic model with only a single fecundity parameter *ω_m_* = *ω_f_*, assumed to be equal in males and females and multiplicative (*ω_b_* = *ω_m_ω_f_*), also produced a very poor fit to the observed data, emphasizing the need for more complex models for understanding the fitness costs of the yellow phenotype.

Finally, we sought to identify a minimal model of low complexity, based on a limited number of biological assumptions, which can still describe the observed dynamics reasonably well. Motivated by the observation that the *β* parameter was often estimated to be one, we assumed for this minimal model that sexual selection acts only by lowering the mating success of yellow phenotype males when wild-type females choose their mate. In addition, we assumed that yellow phenotype individuals have either reduced viability (minimal model 1), or reduced fecundity (minimal model 2), with equal costs between the sexes and multiplicative costs in the case of fecundity. Both of these minimal models have only three free parameters (including *N_e_*), yet yielded better AICc values than all other models. In fact, our minimal model 1 yielded the best AICc value of all models tested, suggesting that the data can be explained well with this simple model in which wild-type females show severe mating-bias against males of yellow phenotype 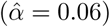 and both sexes with the yellow phenotype experience a modest reduction in viability 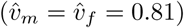. Note, however, that the inclusion of preferential mate choice is crucial in this model, given that the model with only viability selection produced the worst fit among all models tested.

One key advantage of a ML-based approaches is that confidence intervals of parameter estimates can be easily calculated by likelihood ratio tests. Figure 4A demonstrates this for the effective population size parameter *N_e_* in our full model, which we inferred to be *N_e_* = 167 with a 95% confidence interval of approximately 100 − 250, or about 5-12% of the census population size. This estimate is consistent with previous cage evolution experiments in *D. melanogaster*, where *N_e_* values were observed ranging from 4%- 25% of population census sizes. ^28, 31^ Importantly, in our framework, the inference parameter *N_e_* will also depend on the overall goodness-of-fit of the inference model. A misspecified model would be expected to result in lower *N_e_* estimates, because the ML approach would have to assume higher amounts of drift to explain the larger deviations between the predicted and observed frequency trajectories.

**Figure 4:**
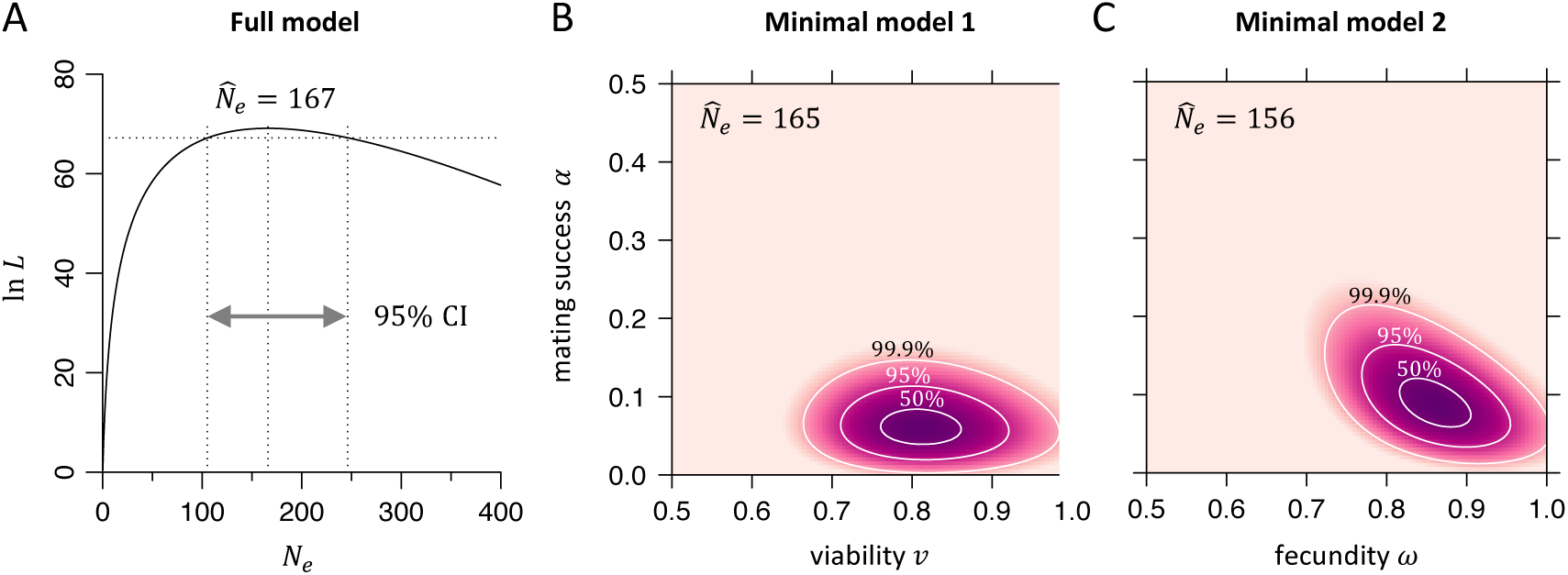
Log-likelihood surfaces and confidence intervals. (A) The log-likelihood curve of our full model varying *N_e_*, while keeping other parameters fixed at at their maximum likelihood estimate. The vertical dashed lines indicate the maximum likelihood estimate 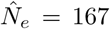 and the corresponding 95% confidence interval (CI), estimated from a likelihood ratio test with one degree of freedom. (B) The log-likelihood surface for minimal model 1 with the mating success parameter *α*, viability parameter *v_m_* = *v_f_*, and fixed 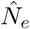. (C) The log-likelihood surface for minimal model 2 with the mating success parameter *α*, fecundity parameter *ω_m_* = *ω_f_*, *ω_b_* = *ω_m_ω_f_*, and fixed *N_e_*. Darker colors correspond to higher likelihoods. The three contour lines specify confidence intervals according to a likelihood ratio test with two degrees of freedom.

Figures 4B,C show the likelihood surface plots and confidence intervals for the selection parameters in each of our two minimal models, demonstrating that our ML approach has good power in inferring these parameters. The diagonal elongation of the contour areas in minimal model 2 suggest that the two parameters are in fact partially dependent on one another in this model.

## Discussion

In this study, we developed a ML approach for estimating selection parameters from time series data of genotype frequencies. The key advancement of our approach over existing methods is the ability to explicitly distinguish among the different components that constitute fitness, including mating success, fecundity, and viability. While our framework can be applied to a wide range of scenarios, we demonstrated it in this study for the specific example of a recessive X-linked allele potentially undergoing sex and phenotype-based sexual, fecundity, and viability selection. Our analysis indicates that these forms of selection indeed can result in distinct genotype frequency trajectories, which can be reliably distinguished from each other using our inference method.

As a proof-of-principle, we applied our approach to study the fitness costs associated with a disrupted *yellow* allele in *D. melanogaster*, using data from cage evolution experiments. Consistent with previous studies,^13, 21^ we found that yellow phenotype males indeed experience a large fitness cost compared to their wild-type counterparts. However, our ML approach allowed us to further show that this primarily results from wild-type females strongly preferring to mate with wild-type males over yellow males. Additionally, yellow phenotype males and females may both experience a moderate reduction in viability according to our minimal model 1 that yielded the highest statistical support (Table 2).

Figure 5 shows that dissecting individual fitness components is crucial for an accurate description of the genotype frequency dynamics at the *yellow* locus. A simplistic model that seeks to approximate this with a single selection parameter would predict significantly different dynamics from those observed in the cages, whereas our minimal model captures the true dynamics with substantially higher accuracy.

**Figure 5:**
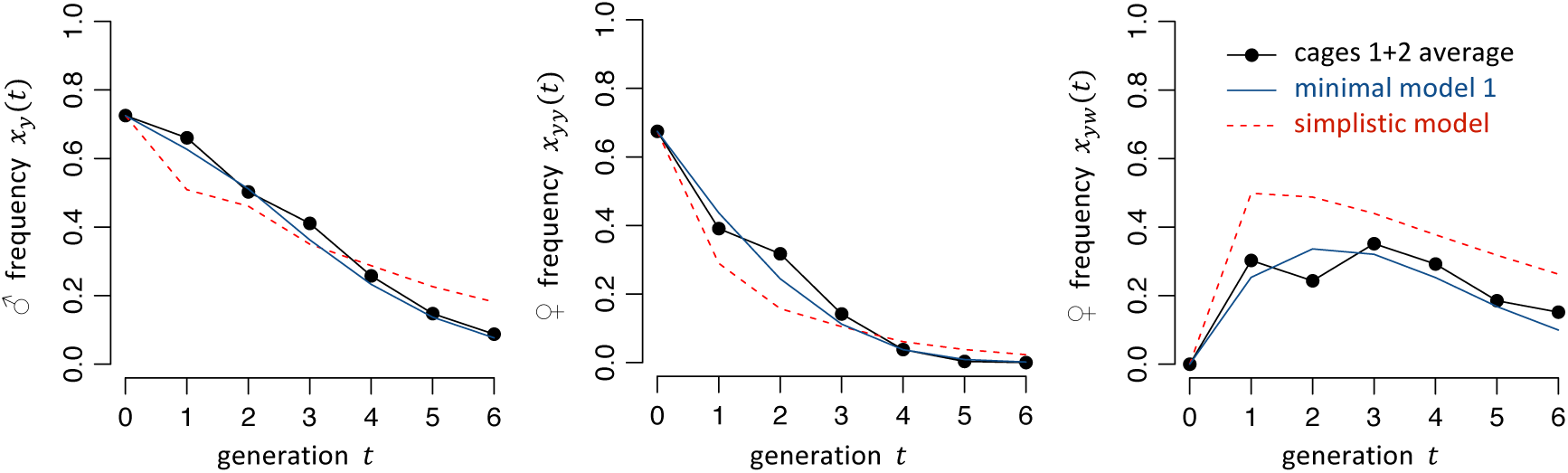
Comparison of the observed genotype trajectories in our population cages (averaged over cages 1 and 2) with the predicted trajectories of the simplistic one-parameter fecundity-selection-only model (*ω_m_* = *ω_f_* = 0.50, *ω_b_* = *ω_m_ω_f_*) and our minimal model 1 (*v_m_* = *v_f_* = 0.81, *α* = 0.06), using the parameter MLEs inferred in Table 2. Simulations were started from the same initial genotype frequencies as the cages and were run without drift.

Despite its generality, there are a number of important limitations to our method. In contrast to many existing methods, which are able to analyze sparse allele frequency data obtained from distant time-points, our method requires frequency estimates over discrete, consecutive generations. This requirement allowed us to establish a fully analytical model based on the Wright-Fisher process, which does not rely on simulations or numerical approximations of the transition probabilities across multiple generations, as are typically invoked by other methods. However, it does likely limit the applicability of our approach to experimental studies where discrete generations can be enforced and data can be obtained for each generation.

There are also potential issues regarding overfitting in our model. For example, in our implementation of a mate choice model, we restricted mate choice selection to males, while females could also potentially undergo such selection. Additionally, with the large number of possible parameters, certain combinations of values can create very similar outcomes. It is likely that such parameters can only be distinguished in complex scenarios involving major sex-based differences between these parameters. This is apparent, for instance, when examining sexual vs. fecundity parameters in Table 2. Similarly, even though our minimal models were alike in that both had mate choice selection against yellow males, the first model had a viability cost for yellow phenotype individuals while the second had a fecundity cost. These results may also be influenced by small sample sizes, so it remains unclear whether or how the yellow phenotype induces a modest reduction in viability, fecundity, or perhaps both.

For calculating the probabilities of specific allele frequency changes between consecutive generations in our model, it was necessary to incorporate an effective population size. We defined this parameter to be equal for males and females and constant across generations, yet other valid possibilities exist. For example, the effective population size could be defined as a fixed percentage of the total population for each generational transition, using either the current generation size, the previous generation size, or some combination of both. Furthermore, male and female likelihoods could be combined, whereas we chose to factorize them in our experimental situation because of a skewed sex ratio likely due to experimental factors. Another possibility would be to treat male and female effective population sizes separately, which would also allow for a lower effective proportion of males compared to females. In some scenarios, these considerations may yield a model that is more representative of the level of stochasticity in each generational transition, though we found that most methods yielded similar results in our limited data set.

Disrupted *yellow* alleles are classic genetic markers in *Drosophila* genetics and have long been known to affect the mating success of male flies due to less effective courtship displays. Specifically, wing extension display is known to be defective in yellow males, resulting in their securing only approximately 10% of matings with a wild-type female when in 1:1 competitions with wild-type males,^13^ which is consistent with our results. Interestingly, in long-term yellow phenotype stocks, yellow females were thought to have evolved to be more receptive to their male counterparts,^3^ but it was later determined that females with the yellow phenotype were themselves more receptive to yellow males, irrespective of their genetic background. ^12^ Our study corroborates this result. While it is possible that yellow females do show a modest preference for wild-type males over yellow males, our study lacked the statistical power to determine this.

Many evolutionary scenarios will still be well served by classical models invoking only a single, fixed selection coefficient. However, certain scenarios will likely require more detailed fitness models such as the one we presented here. Perhaps the primary application for such models involve the spread of selfish genetic elements. The potential use of engineered gene drives,^6, 43^ for example, demands accurate and detailed models for assessing their performance in natural populations, which will likely have to be based on cage trials. For instance, while an existing *D. melanogaster* gene drive^8^ that targeted the *yellow* gene would not be able to spread, a multiplexed-gRNA drive^7^ at this locus could be expected to spread in a frequency-dependent fashion in either of our minimal models involving sexual selection, a feature not commonly associated with such homing-type gene drives. Thus, determination of individual fitness components is not only essential for understanding the spread of gene drives, but could potentially inform their design as well.

## Acknowledgements

We would like to thank Yassi Hafezi for helpful advice on the experimental work. This work was supported by startup funds from the College of Agriculture and Life Sciences at Cornell University to P.W.M, the National Institutes of Health award R21AI130635 to J.C., A.G.C., and P.W.M, and the National Institutes of Health award F32AI138476 to J.C.

